# Inferring membrane properties during clathrin-mediated endocytosis using machine learning

**DOI:** 10.1101/2023.01.11.523591

**Authors:** Zhiwei Lin, Zhiping Mao, Rui Ma

## Abstract

Endocytosis is a fundamental cellular process for eukaryotic cells to transport molecules into the cell. To understand the molecular mechanisms behind the process, researchers have obtained abundant biochemical information about the protein dynamics involved in endocytosis via fluorescence microscopy and geometric information about membrane shapes via electron tomography. However, measuring the biophysical information, such as the osmotic pressure and the membrane tension, remains a problem due to the small dimension of the endocytic invagination. In this work, we combine Machine Learning and Helfrich model of the membrane, as well as the dataset of membrane shapes extracted from the electron tomography to infer biophysical information about endocytosis. Our results show that Machine Learning is able to find solutions that both match the experimental profile and fulfill the membrane shape equations. Furthermore, we show that at the early stage of endocytosis, the inferred membrane tension is negative, which implies strong compressive forces acting at the boundary of the endocytic invagination. This method provides a generic framework to extract membrane information from the super-resolution imaging.

**SIGNIFICANCE:** Endocytosis is a fundamental cellular process that has been extensively studied with the help of fluorescence microscopy and electron microscopy. A large amount of data has been accumulated about the protein dynamics and the membrane shapes. In this work, we combine the widely used Helfrich model and experimental data of membrane shapes to infer the physical information about endocytosis, including the membrane tension and the osmotic pressure. Our work not only proves Machine Learning as a power tool is able to solve the complicated membrane shape equations, but also provides novel biological insights about the initiation of endocytosis in yeast cells.

## INTRODUCTION

Clathrin-mediated endocytosis (CME) is an essential process for eukaryotic cells to uptake nutrients, regulate signal transduction, and control the membrane composition (1–6). When CME occurs, a small patch of the plasma membrane is internalized into the cytoplasm to form an endocytic pit, which is later pinched off to form a vesicle. In the past decades, tremendous efforts have been devoted to understanding the molecular mechanisms that drive CME. Fluorescence microscopy is widely used to obtain the dynamic information of the protein concentration during CME (7–9). Electron microscopy serves as a powerful tool to resolve the morphology of the membrane during CME (10–14). Though the advanced imaging technologies have helped accumulate an enormous amount of data, mining out useful information from the data remains insufficient. For instance, the deformation of the membrane during CME is shaped by many factors, including the force generated by the actin polymerization, the curvature induced by clathrin proteins, and the osmotic pressure as a result of the solute concentration difference between inside and outside of the cell. Geometric profiles of the membrane obtained via the electron tomography therefore contain abundant information of the mechanical properties of the membrane. However, the analysis of the profile data is often limited to a few geometric features, derived from the original high dimensional profile data. How to use the data to have a mechanistic understanding of the physical mechanism remains elusive.

Measuring the physical quantities involved in endocytosis is important for us to understand how endocytosis happens under the orchestrated action of many proteins. Yeast cells have been used as a model system to study endocytosis due to their fast proliferation and easy genetic manipulation (15–17). Different from the mammalian cells, yeast cells have a cell wall, which enables them to maintain a high osmotic pressure (Fig. 1a). Minc et al. have used the deformation of a PDMS chamber to infer the osmotic pressure by assuming the growth rate of fission yeast cells is powered by the osmotic pressure, and the estimated value is 0.85 ± 0.15 MPa (18). Atilgan et al. have obtained a value of 1.5 ± 0.2 MPa in fission yeast cells by comparing the geometric difference between the natural state in which the osmotic pressure expands the cell wall and the lysed state in which the cell wall shrinks in response to the released osmotic pressure (19). In budding yeast, the osmotic pressure is estimated to be 0.6 ± 0.2 MPa by analyzing the volume change upon variations in osmolarity (20). Membrane tension is another important quantity that is relevant for membrane morphology (21–23). Its value might vary significantly among different organisms and is widely accepted to be in the range of 10^−2^ to 10^1^ pN/nm (18, 19, 24–26).

**Figure 1:**
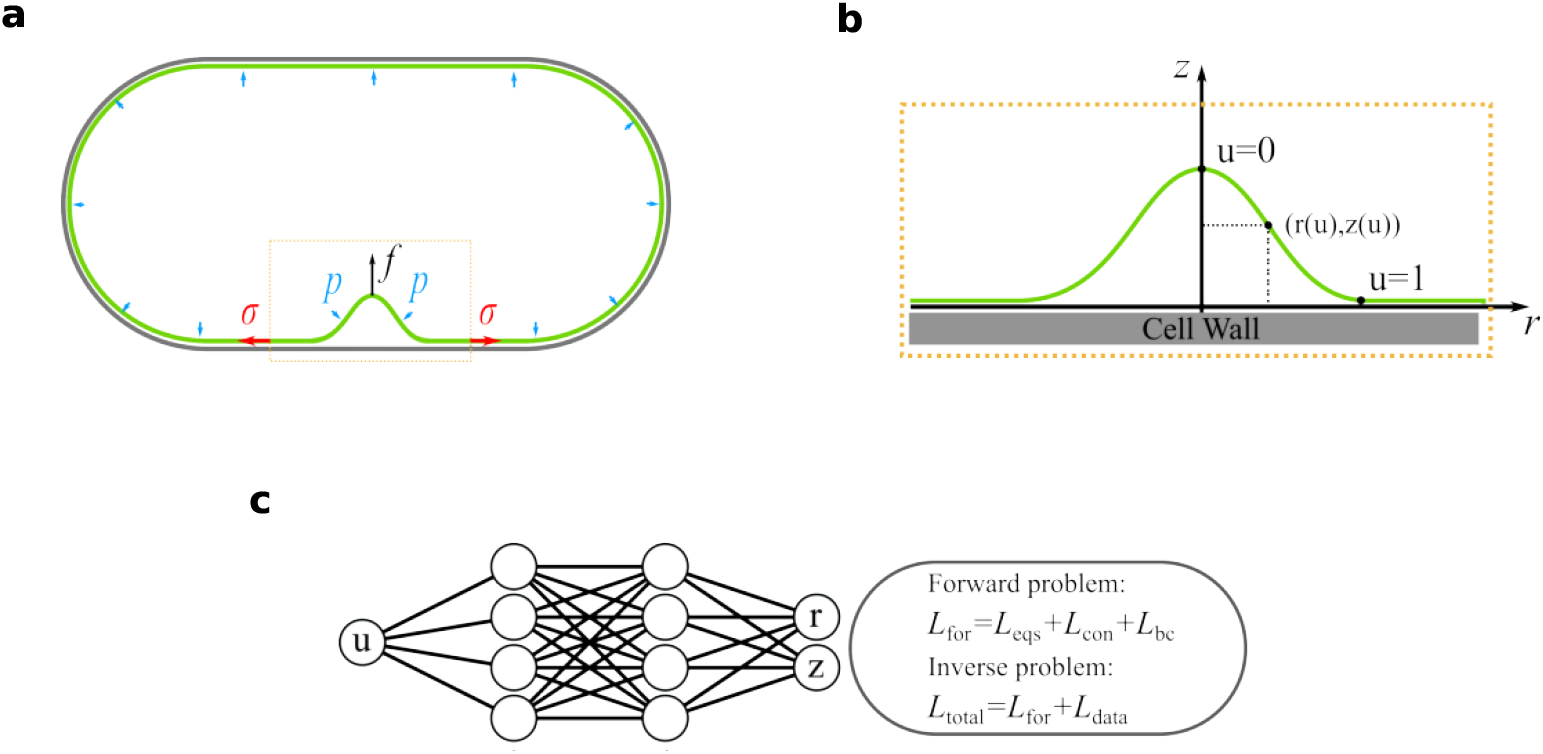
Illustration of the physical parameters of endocytosis and the PINNs method. (a) Simplified illustration of endocytosis in yeast cells. The osmotic pressure inside of the cell pushes the plasma membrane against the cell wall. Internalization of the membrane therefore needs a pulling force *f* to overcome the resistance from the pressure *p*, as well as the membrane tension *σ*. (b) Parametrization of the axisymmetric membrane shape [*r* (*u*), *z*(*u*)] where *u* = 0 corresponds to the membrane tip and *u* = 1 to the membrane base where the membrane is in contact with the cell wall. (c) The structure of the neuron network and the loss functions for the forward and the inverse problem.

Various theoretical models have been proposed to account for the membrane shape evolution during endocytosis (27–32). All of these models are based on the classical Helfrich theory (33, 34), which calculates the membrane shape by minimizing the total energy of the membrane. Physical parameters, such as the bending rigidity, membrane tension, and osmotic pressure are needed to characterize the membrane property. Among the theoretical studies, some assume a very small osmotic pressure and the membrane shape is mainly determined by the membrane tension(28, 30). While other models assume a large osmotic pressure(29, 35). The ambiguity in the choice of these physical parameters might come from the difference between organisms, but also reflects the difficulty in obtaining these physical parameters from experiments.

In the past ten years, Machine Learning (ML) (36, 37) has made brilliant achievements, including pattern recognition (38, 39), computer vision (40, 41), data mining (42, 43), natural language processing (44, 45) and automatic driving (46). Recently, as one of the ML applications in the field of scientific computing, physics-informed neural networks (PINNs) developed in Ref. (47) have been showed to be a powerful tool in solving forward and inverse problems of partial differential equations (PDEs) (48–51). In the framework of PINNs, both the data and the physics contribute to the loss in the training process. PINNs is easy coding and much more flexible than the classical numerical methods. In particular, it has been showed that PINNs is more effective than the classical numerical methods in solving inverse problems, for instance, learning parameterized PDEs (48, 52), and using the concentration field to learn velocity fields with hidden fluid dynamics (52–54).

In this paper we combine the Helfrich model of membrane and PINNs to learn the model parameters including osmotic pressure and membrane tension based on the membrane profile data obtained via the electron tomography for yeast cells (10). The method demonstrates stable convergences of the estimated parameters and, the learned osmotic pressure is consistent with experimental measurements. Furthermore, we find negative membrane tensions at the early stage of endocytosis, which implicates strong compressive forces applied at the boundary of the endocytic patch.

## METHODS

### Helfrich Model of the membrane

We use the classical Helfrich model (34) to calculate the shape of the membrane. The total energy *E* of the membrane is written as

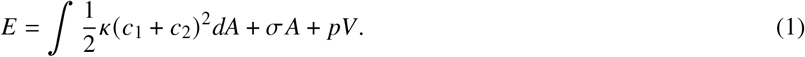

The first term describes the bending energy of the internalized membrane patch with a bending rigidity *κ*. The two principal curvatures of the membrane surface are denoted as *c*_1_ and *c*_2_. The second term describes the energy contribution from the membrane tension *σ* with the conjugated variable *A* being the total surface area of the internalized membrane patch. The third term describes the energy contribution from the osmotic pressure *p* with the conjugated variable *V* being the volume enclosed between the membrane patch and the cell wall (Fig. 1a).

We assume rotational symmetry of the membrane shape such that the surface of the membrane can be parameterized as

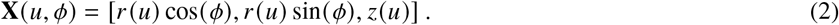

The parameter *ϕ* ∈ [0, 2*π*] is the azimuthal angle, and the parameter *u* ∈ [0, 1] is the meridional coordinate with *u* = 0 at the membrane tip and *u* = 1 at the membrane base where the membrane is in contact with the cell wall. The functions *r* (*u*) and *z*(*u*) depict the membrane profile as shown in Fig. 1b. The total energy (1) then becomes a functional of *r* (*u*), *z*(*u*) and their derivatives up to second order,

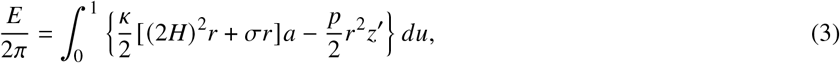

in which

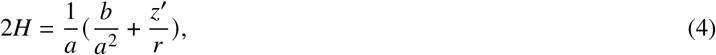

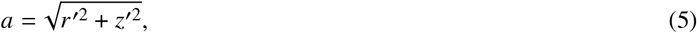

and

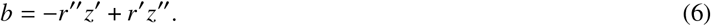

By performing variations against the functions *r* (*u*) and *z*(*u*), with a constraint that *a*(*u*) is a constant, we obtain two variational equations

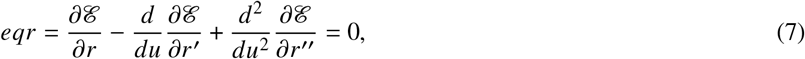

and

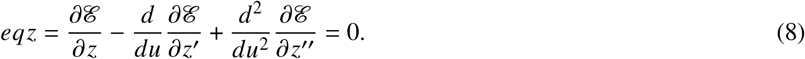

Here we do not give the explicit expressions of the equations which are quite lengthy and contain little information. Note that Eqs. (7) and (8) are fourth-order ordinary differential equations about *r* (*u*) and *z*(*u*). In order to solve the equations, we also need to specify 8 boundary conditions, which include setting the base radius *r* (1) = *R*_*b*_, and the membrane height *z*(0) = *z*_0_. A more detailed description of the boundary conditions can be found in Supplementary information.

### PINNs as a tool to estimate the model parameters

In the forward problem, we solve the membrane shape equations (7) and (8) for a given set of model parameters *κ*, *σ*, *p* and boundary conditions. To do so, we construct a PINN, which is a fully connected neuron network, with the parameter *u* ∈ [0, 1] as the input, and the meridian coordinates of the membrane profile *r* and *z* as the output (Fig. 1 c). The loss function of the network *L*_for_ = *L*_eqs_ + *L*_bc_ + *L*_con_ includes evaluations of the variational equations on a number of points *L*_eqs_, boundary conditions *L*_bc_ that fix the membrane height *z*(0) = *z*_0_ and base radius *r* (1) = *R*_*b*_, as well as coordinate constraints *L*_con_.

In the inverse problem, the model parameters become internal variables of the neuron network and are inferred by the network by comparing with the experimental data. The loss function *L*_tot_ = *L*_for_ + *L*_data_ incorporates the difference between the symmetrized experimental profile and the ML outputs *L*_data_ (Fig. 1c). Note that we assume axisymmetry of the membrane profile in the Helfrich model, therefore perform a symmetrization procedure to the experimental profile for comparison. A detailed description of the neuron network and the symmetrization procedure can be found in Supplementary information.

After learning the model parameters, we further calculate the parameter *f*, which is a Lagrangian multiplier derived from the learned *p*, *σ*, *R*_b_ to impose the membrane height *z*_0_. It represents the minimum force needed to pull the membrane to the corresponding height *z*_0_. A detailed description of the force calculation can be found in Supplementary information.

## RESULTS

### PINNs are able to solve the nonlinear membrane shape equations

We first verify whether PINNs can solve the fourth-order nonlinear membrane shape equations (7) and (8) provided the parameters are given, i.e., the forward problem, by comparing the ML solution that minimizes the loss function *L*_for_ with the solution obtained by a finite difference (FD) method.

The parameters *p*/*κ* and *σ*/*κ* define two characteristic length scales 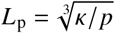 and 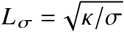. We choose two sets of parameters (listed in Table 1) with one having *L*_p_ < *L* _*σ*_ (pressure-dominant) and the other *L*_p_ > *L* _*σ*_ (tension-dominant). Using a fully connected network with 3 hidden layers, each layer containing 32 nodes, we see an excellent agreement between the ML solutions and the FD solutions for a series of membrane heights for both sets of parameters (Fig. 2). We stress that though the ML solutions are comparable with the FD solutions in accuracy, the ML method is much more time-consuming than the FD method when solving the forward problem. The advantage of the ML method compared with the FD method is mainly reflected in the inverse problem, which is discussed in the next section.

**Table 1:** Two sets of parameters used in Fig. 2.

| $\kappa$ | $p$ | $\sigma$ | $R_b^a$ | $z_0^b$ |
| --- | --- | --- | --- | --- |
| $20 k_B T$ | 1 kPa | 0.01 pN/nm | 50 nm | 10/30/50/100 nm |
| $2000 k_B T$ | 1 MPa | 0.01 pN/nm | 50 nm | 10/30/50/100 nm |

<sup>a</sup> $R_b = r(1)$ , referred to the base radius of the membrane.
<sup>b</sup> $z_0 = z(0)$ , referred to the height of the membrane.

**Figure 2:**
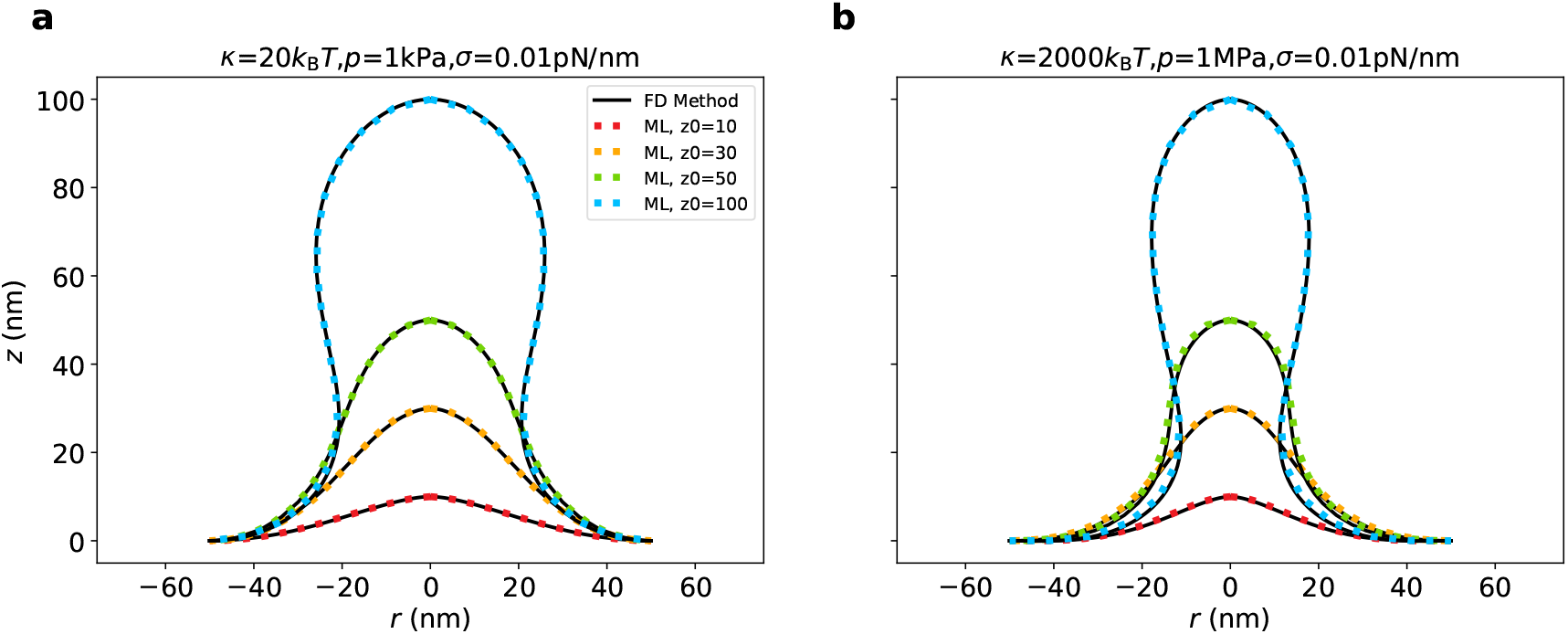
The ML solutions are comparable with the FD solutions in accuracy. (a, b) ML solutions (dashed lines) for a series of membrane heights *z*_0_ = 10, 30, 50, 100 nm are shown in different colors. The corresponding FD solutions are shown in solid lines. The tension-dominant regime is shown in (a) and the pressure-dominant regime is shown in (b). The two sets of parameters are listed in Table 1.

### PINNs show stable convergence of the model parameters for the inverse problem

In this section, we use the PINNs to infer the model parameters, which include the re-scaled pressure *p*/*κ*, the re-scaled tension *σ*/*κ*, and the base radius *R*_b_. The network parameters, which contain the model parameters, are trained to minimize the total loss function *L*_tot_. In this way, the ML outputs not only fulfil the variational equations, but also match the membrane profile data obtained via electron tomography. We stress that it is the re-scaled parameters *p*/*κ* and *σ*/*κ* that can be inferred from the experimental data. In order to have the absolute value of the membrane tension *σ* and the osmotic pressure *p*, we specify *κ* = 2000 *k*_B_*T* from Ref. (29) for the rest of the paper.

During the training of the neural network, we switch the model parameters between trainable and untrainable states. In particular, we fix *σ*/*κ* and *p*/*κ* and tune *R*_b_ in the first 10^5^ training epochs, then fix *R*_b_ and vary *σ*/*κ* and *p*/*κ* for another 10^5^ training epochs. After executing this training procedure twice, the loss function *L*_tot_ and *L*_data_ can be both reduced to 10^−2^ (Fig. 3a), which implies that the membrane shape equations are satisfied and the ML-learned shape fits well with the experimental data. All three parameters show a trend of convergence at the end of the training (Fig. 3b). Upon switching of the training states, their values show a jump with the drop of the loss function (Fig. 3b).

**Figure 3:**
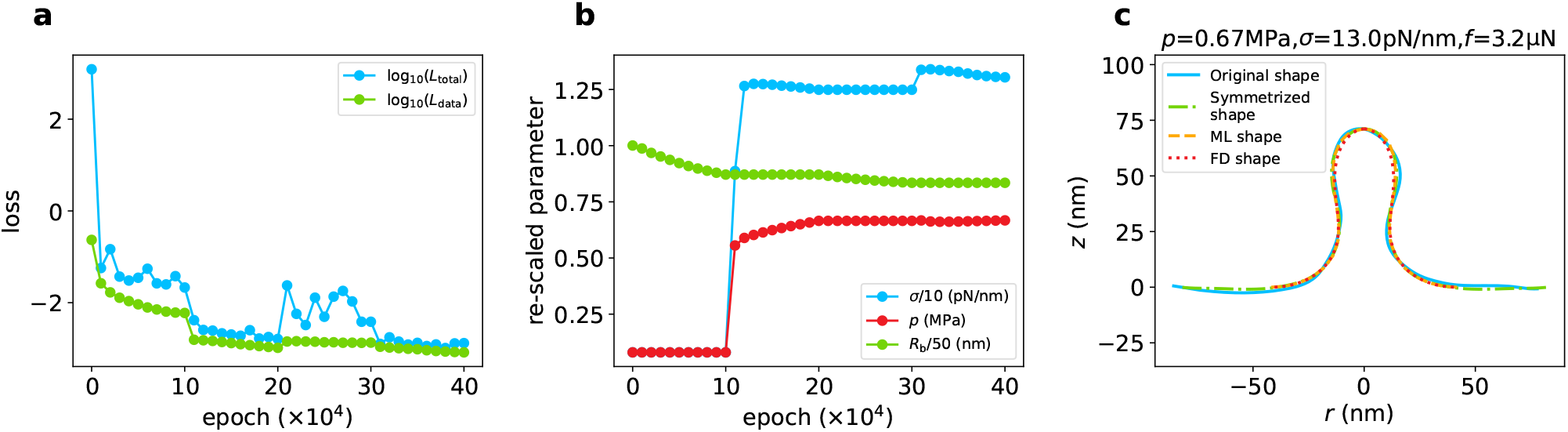
Convergence test of the PINNs and effectiveness test of the ML-predicted parameters. (a) Evolution of the total loss function *L*_tot_ and the experimental loss function *L*_data_ during a training process. (b) Evolution of the three model parameters to be tuned by the neuron network to minimize the total loss function during a training process. (c) Illustration of the original membrane profile (solid cyan), the symmetrized membrane profile (dash-dotted green), the ML-predicted membrane profile (dashed orange), and the FD-solved membrane profile, respectively.

As a verification of the effectiveness of the ML-learned parameters, we substitute them into the membrane shape equations and solve the equations with a FD method. We overlay the original and symmetrized experimental membrane profile, the ML-learned membrane profile, the FD-solved membrane profile, and observe a good agreement between all the profiles (Fig. 3c).

Due to the non-linearity of the membrane shape equations (7) and (8), the solutions might depend on the model parameters with different sensitivity. The possibility that different parameters lead to similar membrane shapes limits the precision of the estimated parameters. In order to estimate the precision of the parameters for a particular experimental profile, we repeat the learning procedure 10 times and use the standard deviation of the parameters over the 10 times to measure the uncertainty of the estimation. For most of the experimental profiles, the multiple learning strategy achieves a small standard deviation compared with the average value. In particular, for the profile shown in Fig. 3c, the learned osmotic pressure *p* = 0.71 ± 0.06 MPa, and the membrane tension *σ* = 13.7 ± 0.18 pN/nm (see the comparison between the ML-learned shape and the experimental profile for the other 9 trainings in Fig. S2).

### ML-predicted parameters are consistent with experimental measurements

We perform the ML learning procedure on 79 membrane profiles extracted from the Time-Resolved Electron Tomography provided by Ref. (10). For each dataset, the learning procedure was repeated 10 times, so that we have 790 learning results in total for the osmotic pressure *p* and the membrane tension *σ*. In most of the learnings, the total loss function can be reduced to below 10^−2^ and concentrated at 10^−3^ (Fig. 4), which proves the effectiveness of the ML method. A more direct visualization of the one-time learning results for 9 randomly picked-up membrane profiles are shown in Fig. 5. All of the learnings show good agreement among the ML-learned shape, the symmetrized experimental profile, and the FD-solved shape, which proves the effectiveness of the ML method. We stress that what learned by the ML method is the symmetrized experimental profile, but not the original profile. For original profiles that are highly asymmetric, the membrane might be subject to asymmetric force distributions or local variations of the model parameters. The ML-learned parameters therefore represent the average of the model parameters such that the local variations are evened out.

**Figure 4:**
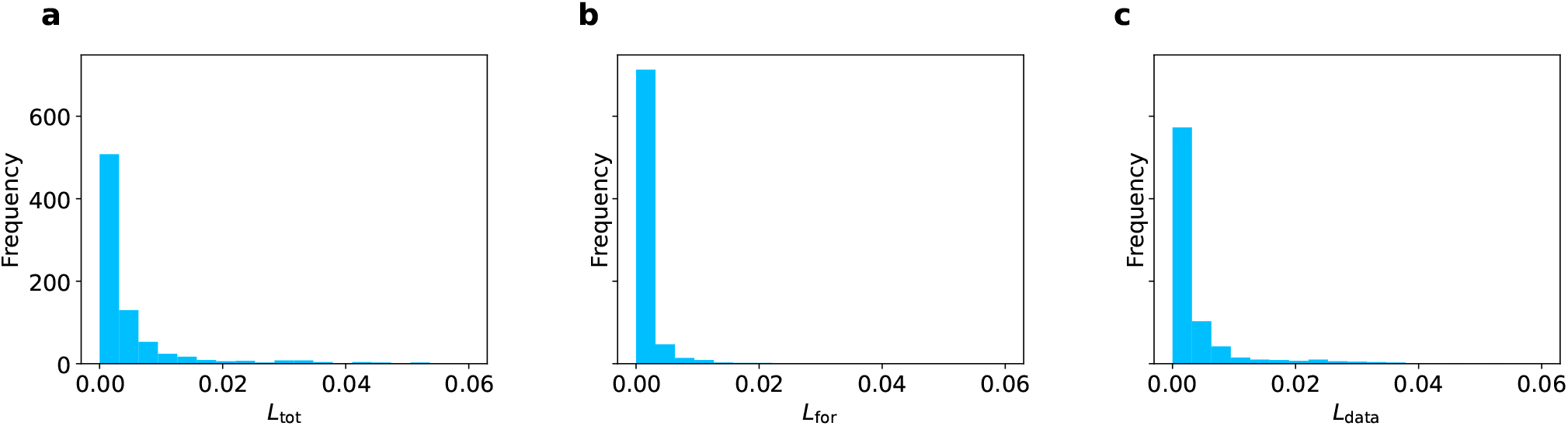
Histogram of the trained loss functions for 79 experimental profiles. Histograms of the loss function *L*_tot_ in (a), *L*_for_ in (b) and *L*_data_ in (c).

**Figure 5:**
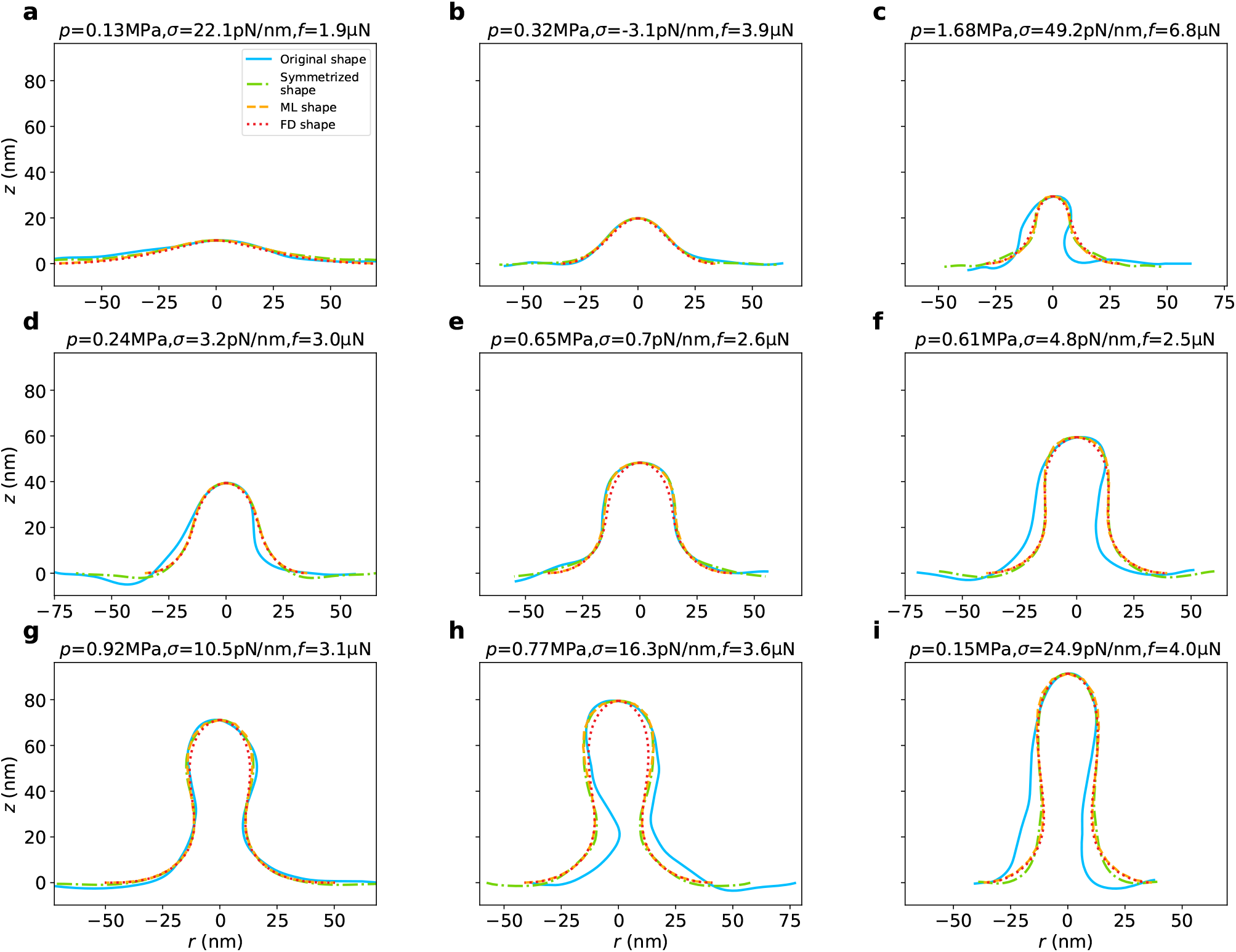
Illustration of the learned results for 9 experimental membrane profiles of various heights. The learned parameters *σ*, *p* and the derived parameter *f* are indicated on the top of each box. In each panel, the original membrane profile (solid cyan), the symmetrized membrane profile (dash-dotted green), the ML-learned membrane profile (dashed orange), and the FD-solved membrane profile (dotted red), are shown respectively.

To demonstrate the precision of learned parameters, for each one of the 79 experimental profiles, we calculate the average and the standard deviation of the learned parameters over the 10 repetitions and show their joint distribution in Fig. 6. We find that: (i) The distribution of the standard deviations is concentrated in a small range near 0, and is much narrower than the distribution of the average values. This certifies a good precision of the learned parameters; (ii) The average values of the osmotic pressure *p* are mostly positive and the distribution is peaked at 0.45 MPa, which is consistent with experimental measurements (18, 19); (iii) The distribution of the average values of the membrane tension *σ* exhibits two peaks, a large one centered at −100 pN/nm and a small one centered at 12 pN/nm; (iv) The average values of the force *f* are mostly positive and peaked at 2.8 *μ*N.

**Figure 6:**
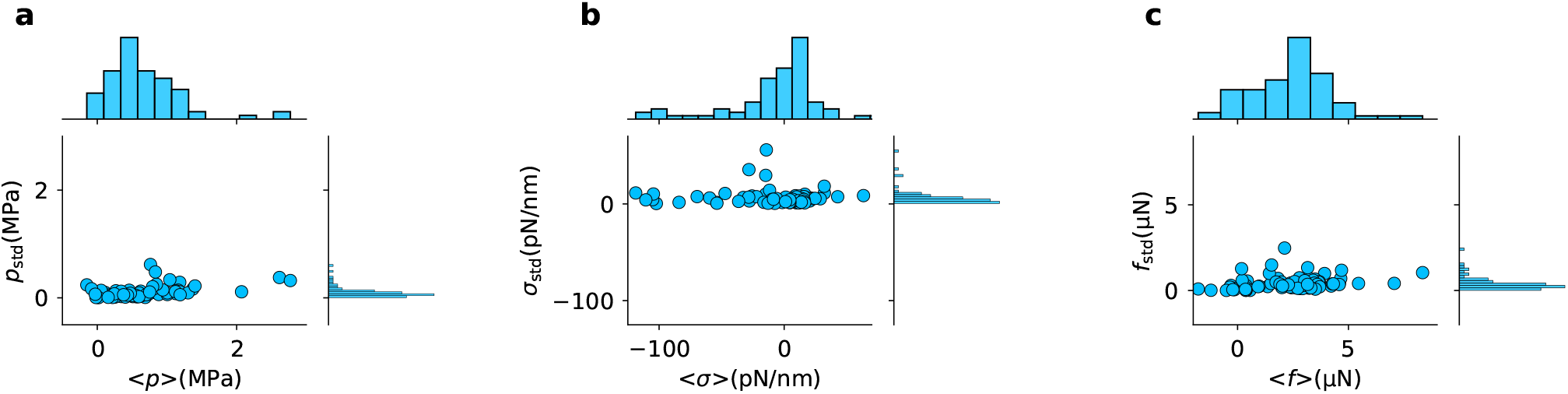
Joint distribution of the average and the standard deviation for ML-learned parameters. The distribution for the osmotic pressure *p* in (a), for the membrane tension *σ* in (b), and for the force *f* in (c).

### Negative membrane tensions occur at early stage of endocytosis

We have shown that the average values of the model parameters have a much wider distribution than the standard deviations over the 10 repeated learnings (Fig. 6), which suggests that the parameters might vary at different stages of endocytosis. To test the stage-dependence of the model parameters, we use the membrane height as a indicator of the timeline of endocytosis, and plot the ML-learned parameters as a function of the corresponding membrane height *z*_0_. It is found that the osmotic pressure *p* shows no dependence on the membrane height (Fig. 7a), but the membrane tension *σ* exhibits a strong height dependence (Fig. 7b). Large and negative membrane tensions are found for *z*_0_ < 25 nm. Above 50 nm, the membrane tensions are almost independent of the membrane height *z*_0_ and remain to be small and positive (Fig. 7b). The forces *f* stay around 0 *μ*N for *z*_0_ < 25 nm, and reach a plateau of about 3 *μ*N above 50 nm (Fig. 7c). A strong positive correlation (*R* = 0.87) between the force *f* and the membrane tension *σ* is observed (Fig. 7d). However, the force *f* is only weakly correlated with the osmotic pressure *p* (*R* = 0.25)(Fig. 7e).

**Figure 7:**
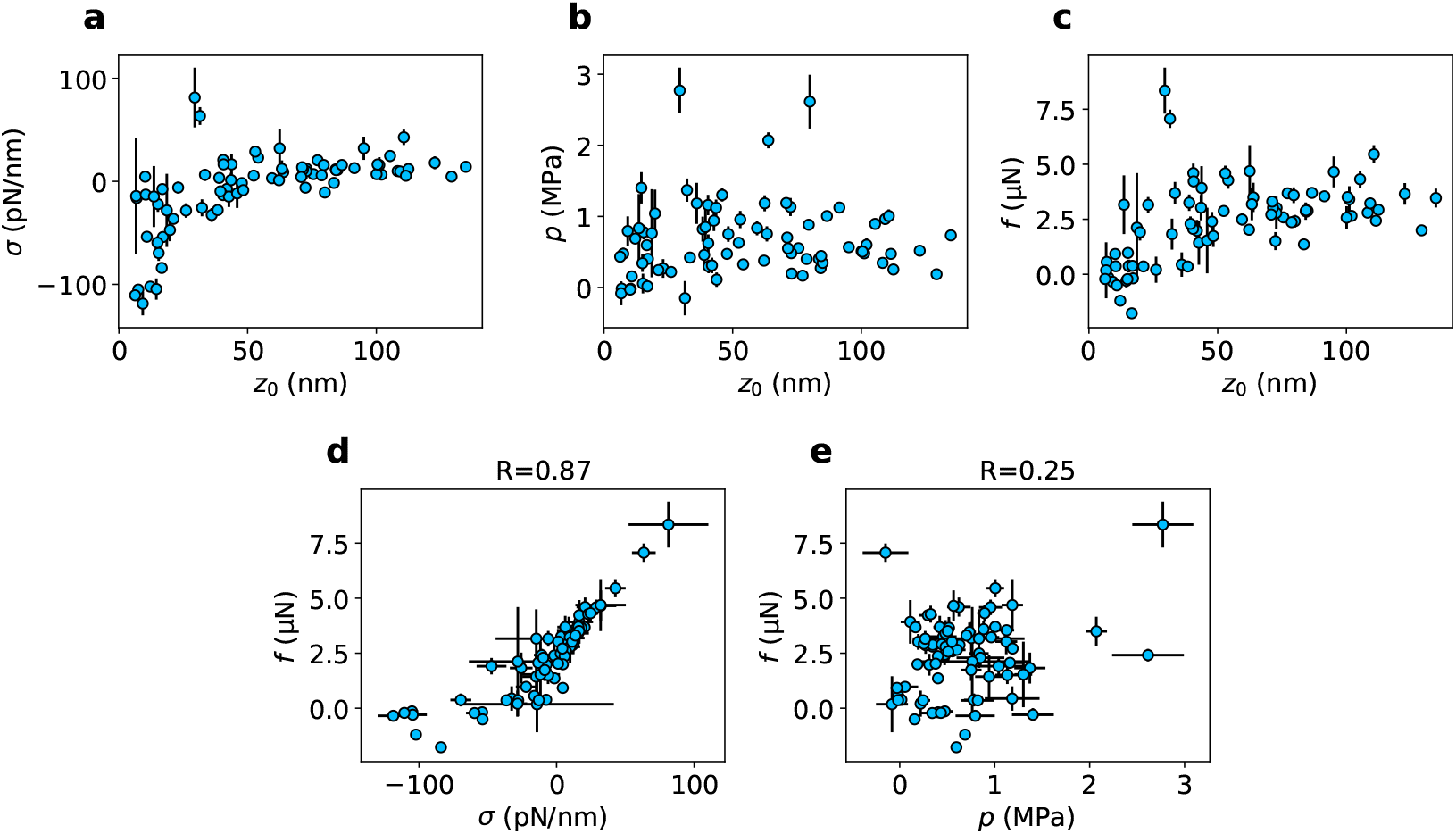
Statistical analysis of ML-learned parameters. (a-c) Model parameters as a function of the membrane height, with the the osmotic pressure *p* in (a), the membrane tension *σ* in (b), and the force *f* in (c). Each point represents the average value of the same experimental profile over 10 repeated learnings, and the error bar represents the standard deviation. (d,e) Scattered plots of (*f*, *σ*) in left, and (*f*, *p*) in right. The horizontal and vertical error bars of each point represent the standard deviations of the corresponding parameter over 10 repeated learnings, respectively.

## DISCUSSION

### The ML method has the natural advantage in estimating model parameters

We have shown that the ML method is able to solve the nonlinear membrane shape equations to an accuracy that is comparable with the FD method. However, in terms of speed, the FD method beats the ML methods with orders of magnitude. The ML method takes at least a few minutes to solve the equations, and even hours when the membrane height is large. In contrast, it takes less than a second for the FD method to solve the same equation. However, when it comes to the inverse problem, i.e., learning the model parameters with given experimental data, the FD method is less flexible than the ML method. We use the bvp5c solver in MATLAB which is an iterative algorithm based on the finite difference scheme of Runge-Kutta methods (55). It requires a proper initial guess of the solutions to solve the membrane shape equations provided the parameters are given. In practice, we always choose a flat shape as the initial guess. If the membrane height is large, direct use of the bvp5c solver often causes error. By contrast, the ML method does not require a proper guess of the solution, and is naturally extended to solve the inverse problem of the membrane shape equations by incorporating *L*_data_ into the loss function. Solving the inverse problem with the ML method has almost the same computational cost as the forward problem. Therefore, the ML method is very suitable for the task of parameter learning.

### Our results implies that the initiation of endocytosis is facilitated with negative membrane tension

The presence of a large osmotic pressure inside of the yeast cells imposes a large force barrier to complete endocytosis (35). In this paper, we have shown that the ML-learned osmotic pressure has an average value of about 0.66 MPa (Fig. 7a), and the force *f* to pull the membrane to a height of 100nm is about 3 *μ*N (Fig. 7c). These findings are consistent with previous studies. However, at the early stage of endocytosis when the membrane height is low, our results suggest large and negative membrane tensions are present at the boundary of the endocytic invagination (Fig. 7b and c). The negative tension implies an inward force to pump lipids into the endocytic invagination through the boundary and might induce membrane buckling, thus reducing the force requirement to pull the membrane up. The myosin-1 motors have been reported to form a ring structure around the endocytic invagination (56). They might play such a role to generate negative tensions. Investigating the effect of the negative tension will be our future work.

### Sensitivity of the model parameters limit the precision of the ML estimation

There are two possible factors that limit the precision of the learned parameters. One is due to the neuron network being unable to converge in a limited number of iterations, which is manifested as a large loss function. The other is due to the fact that different model parameters could give similar membrane shapes. In our results, we find for some experimental profiles, the repeated learnings indeed give quite different parameters. The large error bars observed in some of the points in Fig. 7 mainly arise from the insensitivity of the model parameters to the membrane shapes but not the failure of the neuron network. To prove this, we overlay the FD-solved membrane shapes of the 10 repeated learnings for experimental profiles that have the largest error bars. Most of the shapes overlap with each other, though the parameters are quite different (Fig. 8). In addition, for the 79 experimental profiles, we observe no correlation between the average loss function and the standard deviation of the model parameters over the 10 repeated learnings (Fig. 9), which implies that the large error bars are not due to the failure of the neuron network. Improving the parameter precision for those profiles therefore needs more information than the geometric shapes.

**Figure 8:**
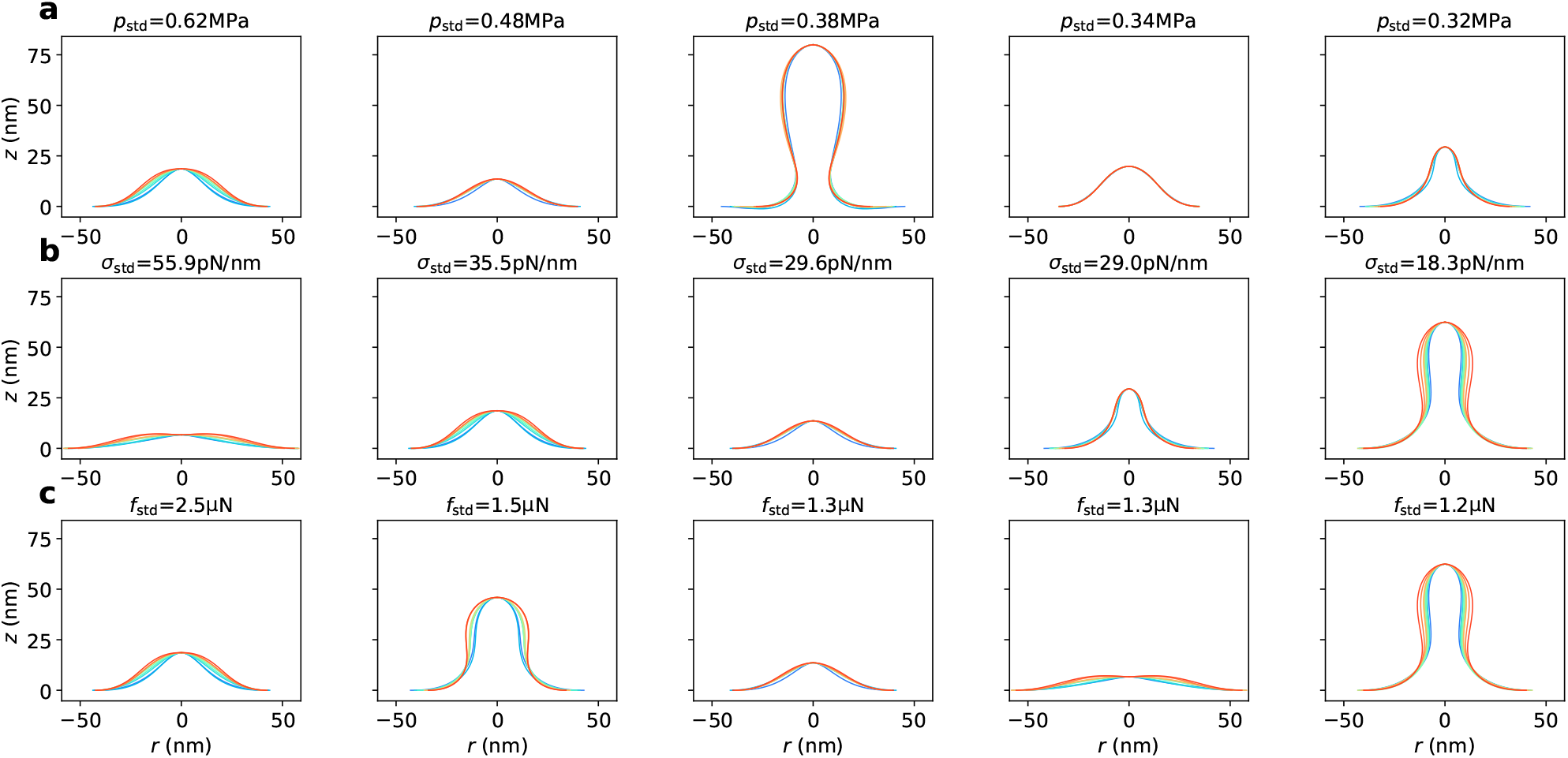
Learned membrane shapes for experimental profiles with the largest standard deviations of model parameters. Each panel shows the FD-solved membrane shapes for the same experimental profile over 10 repeated learnings. The top, middle and bottom rows are for parameters *p*, *σ* and *f*, respectively. From left to right, the standard deviations go from large to small.

**Figure 9:**
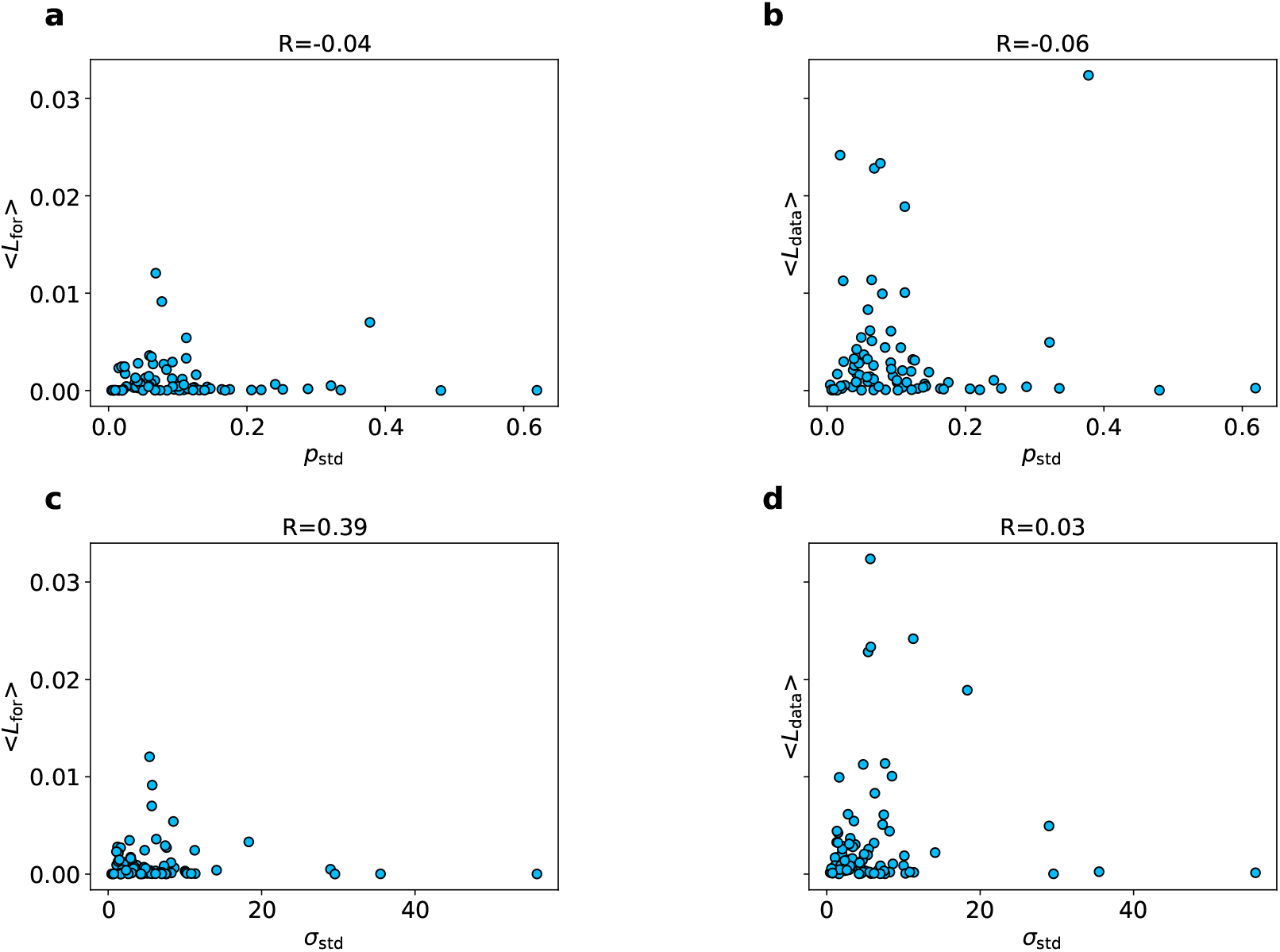
Scattered plots of the standard deviations and the average loss functions over 10 repeated learnings. (a, b) The average loss functions ⟨*L*_for_⟩ vs. the standard deviation of the osmotic pressure *p*_std_ in (a) and ⟨*L*_data_⟩ vs. *p*_std_ in (b). (c, d) The average loss functions ⟨*L*_for_⟩ vs. the standard deviation of the membrane tension *σ*_std_ in (a) and ⟨*L*_data_⟩ vs. *σ*_std_ in (b).

## CONCLUSION

In this paper, we combine the ML method with the experimental data on membrane profiles obtained via electron tomography to infer the mechanical parameters involved in endocytosis in yeast cells. We show that the ML method is able to solve the nonlinear membrane shape equations to an accuracy that is almost indistinguishable from the traditional FD method, and demonstrates advantages over the FD method in estimating the model parameters from the existing data. The estimated osmotic pressure has an average value of 0.66 MPa, which is slightly smaller that the direct measurement results (0.85 MPa in Ref. (18) and 1.5 MPa in Ref. (19)). In addition, the ML method predicts a very negative membrane tension for low membrane heights, which suggests the presence of compressive boundary forces at the base of the membrane. These forces might help initiation of the endocytosis by reducing the forces needed to pull the membrane up against the high osmotic pressure.

## AUTHOR CONTRIBUTIONS

R.M. conceived the project; R.M. and Z.M. designed the study and Z.L. performed the computational work; R.M. and Z.M. supervised the project; Z.L., R.M. and Z.M. wrote the paper.

## ADDITIONAL INFORMATION

Supplementary Information accompanies this paper at “Supplementary information.pdf”.

## DATA AVAILABILITY

Database of membrane profiles obtained by electron microscopy are available in https://www.embl.de/download/briggs/endocytosis.html.

## CODE AVAILABILITY

The custom code generated during the current study are available at GitHub.

## ACKNOWLEDGMENTS

We thank Prof. Julien Berro for critical reading of the manuscript. RM acknowledges financial support from National Natural Science Foundation of China under Grants No. 12004317, Fundamental Research Funds for Central Universities of China under Grant No. 20720200072, and 111 project No. B16029. ZM acknowledges financial support from National Natural Science Foundation of China under Grants No. 12171404, Fundamental Research Funds for Central Universities of China under Grant No. 20720210037.

## Notes

### Competing Interest Statement

The authors have declared no competing interest.

### Summary of Updates

Figures are revised for cosmetic reasons.

## REFERENCES

1. McMahon, H. T., and E. Boucrot, 2011. Molecular mechanism and physiological functions of clathrin-mediated endocytosis. Nature reviews Molecular cell biology 12:517–533.

2. Sorkin, A., and M. A. Puthenveedu, 2013. Clathrin-mediated endocytosis. *In* Vesicle Trafficking in Cancer, Springer, 1–31.

3. Lu, R., D. G. Drubin, and Y. Sun, 2016. Clathrin-mediated endocytosis in budding yeast at a glance. Journal of Cell Science 129:1531–1536.

4. Kaksonen, M., and A. Roux, 2018. Mechanisms of clathrin-mediated endocytosis. Nature reviews Molecular cell biology 19:313–326.

5. Lacy, M. M., R. Ma, N. G. Ravindra, and J. Berro, 2018. Molecular mechanisms of force production in clathrin-mediated endocytosis. FEBS letters 592:3586–3605.

6. Mettlen, M., P.-H. Chen, S. Srinivasan, G. Danuser, and S. L. Schmid, 2018. Regulation of clathrin-mediated endocytosis. Annual review of biochemistry 87:871.

7. Coffman, V. C., and J.-Q. Wu, 2012. Counting protein molecules using quantitative fluorescence microscopy. Trends in biochemical sciences 37:499–506.

8. Wu, J.-Q., and T. D. Pollard, 2005. Counting cytokinesis proteins globally and locally in fission yeast. Science 310:310–314.

9. Kaksonen, M., C. P. Toret, and D. G. Drubin, 2006. Harnessing actin dynamics for clathrin-mediated endocytosis. Nature reviews Molecular cell biology 7:404–414.

10. Kukulski, W., M. Schorb, M. Kaksonen, and J. A. Briggs, 2012. Plasma membrane reshaping during endocytosis is revealed by time-resolved electron tomography. Cell 150:508–520.

11. Avinoam, O., M. Schorb, C. J. Beese, J. A. Briggs, and M. Kaksonen, 2015. Endocytic sites mature by continuous bending and remodeling of the clathrin coat. Science 348:1369–1372.

12. Sochacki, K. A., A. M. Dickey, M.-P. Strub, and J. W. Taraska, 2017. Endocytic proteins are partitioned at the edge of the clathrin lattice in mammalian cells. Nature cell biology 19:352–361.

13. Sochacki, K. A., and J. W. Taraska, 2019. From flat to curved clathrin: controlling a plastic ratchet. Trends in cell biology 29:241–256.

14. Sochacki, K. A., and J. W. Taraska, 2021. Find your coat: using correlative light and electron microscopy to study intracellular protein coats. Current Opinion in Cell Biology 71:21–28.

15. Low, P. S., and S. Chandra, 1994. Endocytosis in plants. Annual review of plant biology 45:609–631.

16. Aghamohammadzadeh, S., and K. R. Ayscough, 2009. Differential requirements for actin during yeast and mammalian endocytosis. Nature cell biology 11:1039–1042.

17. Basu, R., E. L. Munteanu, and F. Chang, 2014. Role of turgor pressure in endocytosis in fission yeast. Molecular biology of the cell 25:679–687.

18. Minc, N., A. Boudaoud, and F. Chang, 2009. Mechanical forces of fission yeast growth. Current Biology 19:1096–1101.

19. Atilgan, E., V. Magidson, A. Khodjakov, and F. Chang, 2015. Morphogenesis of the fission yeast cell through cell wall expansion. Current Biology 25:2150–2157.

20. Schaber, J., M. A. Adrover, E. Eriksson, S. Pelet, E. Petelenz-Kurdziel, D. Klein, F. Posas, M. Goksör, M. Peter, S. Hohmann, et al., 2010. Biophysical properties of Saccharomyces cerevisiae and their relationship with HOG pathway activation. European Biophysics Journal 39:1547–1556.

21. Gustin, M. C., X.-L. Zhou, B. Martinac, and C. Kung, 1988. A mechanosensitive ion channel in the yeast plasma membrane. Science 242:762–765.

22. Boulant, S., C. Kural, J.-C. Zeeh, F. Ubelmann, and T. Kirchhausen, 2011. Actin dynamics counteract membrane tension during clathrin-mediated endocytosis. Nature cell biology 13:1124–1131.

23. Wu, X.-S., S. Elias, H. Liu, J. Heureaux, P. J. Wen, A. P. Liu, M. M. Kozlov, and L.-G. Wu, 2017. Membrane tension inhibits rapid and slow endocytosis in secretory cells. Biophysical journal 113:2406–2414.

24. Shibly, S. U. A., C. Ghatak, M. A. S. Karal, M. Moniruzzaman, and M. Yamazaki, 2016. Experimental estimation of membrane tension induced by osmotic pressure. Biophysical journal 111:2190–2201.

25. Henon, S., G. Lenormand, A. Richert, and F. Gallet, 1999. A new determination of the shear modulus of the human erythrocyte membrane using optical tweezers. Biophysical journal 76:1145–1151.

26. Engelhardt, H., and E. Sackmann, 1988. On the measurement of shear elastic moduli and viscosities of erythrocyte plasma membranes by transient deformation in high frequency electric fields. Biophysical journal 54:495–508.

27. Agrawal, A., and D. J. Steigmann, 2009. Modeling protein-mediated morphology in biomembranes. Biomechanics and modeling in mechanobiology 8:371–379.

28. Walani, N., J. Torres, and A. Agrawal, 2015. Endocytic proteins drive vesicle growth via instability in high membrane tension environment. Proceedings of the national academy of sciences 112:E1423–E1432.

29. Dmitrieff, S., and F. Nédélec, 2015. Membrane mechanics of endocytosis in cells with turgor. PLoS computational biology 11:e1004538.

30. Hassinger, J. E., G. Oster, D. G. Drubin, and P. Rangamani, 2017. Design principles for robust vesiculation in clathrin-mediated endocytosis. Proceedings of the national academy of sciences 114:E1118–E1127.

31. Alimohamadi, H., R. Vasan, J. E. Hassinger, J. C. Stachowiak, and P. Rangamani, 2018. The role of traction in membrane curvature generation. Molecular biology of the cell 29:2024–2035.

32. Napoli, G., and A. Goriely, 2020. Elastocytosis. Journal of the Mechanics and Physics of Solids 145:104133.

33. Jülicher, F., and U. Seifert, 1994. Shape equations for axisymmetric vesicles: a clarification. Physical Review E 49:4728.

34. Helfrich, W., 1973. Elastic properties of lipid bilayers: theory and possible experiments. Zeitschrift für Naturforschung c 28:693–703.

35. Ma, R., and J. Berro, 2021. Endocytosis against high turgor pressure is made easier by partial coating and freely rotating base. Biophysical Journal 120:1625–1640.

36. Mitchell, T. M., and T. M. Mitchell, 1997. Machine learning, volume 1. McGraw-hill New York.

37. Jordan, M. I., and T. M. Mitchell, 2015. Machine learning: Trends, perspectives, and prospects. Science 349:255–260.

38. Theodoridis, S., and K. Koutroumbas, 2006. Pattern recognition. Elsevier.

39. Jain, A. K., R. P. W. Duin, and J. Mao, 2000. Statistical pattern recognition: A review. IEEE Transactions on pattern analysis and machine intelligence 22:4–37.

40. Voulodimos, A., N. Doulamis, A. Doulamis, and E. Protopapadakis, 2018. Deep learning for computer vision: A brief review. Computational intelligence and neuroscience 2018.

41. Forsyth, D. A., and J. Ponce, 2002. Computer vision: a modern approach. prentice hall professional technical reference.

42. Chen, M.-S., J. Han, and P. S. Yu, 1996. Data mining: an overview from a database perspective. IEEE Transactions on Knowledge and data Engineering 8:866–883.

43. Hand, D. J., 2007. Principles of data mining. Drug safety 30:621–622.

44. Chowdhary, K., 2020. Natural language processing. Fundamentals of artificial intelligence 603–649.

45. Manning, C., and H. Schutze, 1999. Foundations of statistical natural language processing. MIT press.

46. Naranjo, J. E., C. González, R. García, T. De Pedro, and R. E. Haber, 2005. Power-steering control architecture for automatic driving. Ieee transactions on intelligent transportation systems 6:406–415.

47. Raissi, M., P. Perdikaris, and G. E. Karniadakis, 2019. Physics-informed neural networks: A deep learning framework for solving forward and inverse problems involving nonlinear partial differential equations. Journal of Computational physics 378:686–707.

48. Lu, L., X. Meng, Z. Mao, and G. E. Karniadakis, 2021. DeepXDE: A deep learning library for solving differential equations. SIAM Review 63:208–228.

49. Karniadakis, G. E., I. G. Kevrekidis, L. Lu, P. Perdikaris, S. Wang, and L. Yang, 2021. Physics-informed machine learning. Nature Reviews Physics 3:422–440.

50. Raissi, M., A. Yazdani, and G. E. Karniadakis, 2020. Hidden fluid mechanics: Learning velocity and pressure fields from flow visualizations. Science 367:1026–1030.

51. Cai, S., Z. Mao, Z. Wang, M. Yin, and G. E. Karniadakis, 2022. Physics-informed neural networks (PINNs) for fluid mechanics: A review. Acta Mechanica Sinica 1–12.

52. Yazdani, A., L. Lu, M. Raissi, and G. E. Karniadakis, 2020. Systems biology informed deep learning for inferring parameters and hidden dynamics. PLoS computational biology 16:e1007575.

53. Mao, Z., A. D. Jagtap, and G. E. Karniadakis, 2020. Physics-informed neural networks for high-speed flows. Computer Methods in Applied Mechanics and Engineering 360:112789.

54. Jagtap, A. D., Z. Mao, N. Adams, and G. E. Karniadakis, 2022. Physics-informed neural networks for inverse problems in supersonic flows. Journal of Computational Physics 466:111402.

55. Kierzenka, J., and L. F. Shampine, 2008. A BVP solver that controls residual and error. JNAIAM J. Numer. Anal. Ind. Appl. Math 3:27–41.

56. Mund, M., J. A. van der Beek, J. Deschamps, S. Dmitrieff, P. Hoess, J. L. Monster, A. Picco, F. Nédélec, M. Kaksonen, and J. Ries, 2018. Systematic nanoscale analysis of endocytosis links efficient vesicle formation to patterned actin nucleation. Cell 174:884–896.

