## Supplemental Materials and Supplemental Figures for "Inferring membrane properties during clathrin-mediated endocytosis using machine learning"

Manuscript submitted to **Biophysical** *Journal*
Article

### Supplementary information for "Inferring membrane properties during clathrin-mediated endocytosis using machine learning"

Zhiwei Lin[1], Zhiping Mao[2,*], and Rui Ma[1,3,*]

[1]Department of Physics, College of Physical Science and Technology, Xiamen University, Xiamen, China
[2]School of Mathematical Sciences, Fujian Provincial Key Laboratory of Mathematical Modeling and High-Performance Scientific Computing, Xiamen University, Xiamen, China
[3]Fujian Provincial Key Laboratory for Soft Functional Materials Research, Research Institute for Biomimetics and Soft Matter, Xiamen University, Xiamen, China

#### VARIATIONAL FORM OF THE MEMBRANE SHAPE EQUATIONS

When using $[r(u), z(u)]$ to describe the membrane profile for $u \in [0,1]$, the free energy can be explicitly expressed as a functional of $r(u)$ and $z(u)$, as well as their derivatives (1),

$$\frac{E}{2\pi} = \int_0^1 \left\{\frac{\kappa}{2}[(2H)^2 r + \sigma r]a - \frac{p}{2}r^2 z'\right\} du, \tag{S1}$$

in which

$$2H = \frac{1}{a}(\frac{b}{a^2} + \frac{z'}{r}), \tag{S2}$$

$$a = \sqrt{r'^2 + z'^2}, \tag{S3}$$

and

$$b = -r''z' + r'z''. \tag{S4}$$

The variational equations then can be written down as

$$eqr[r, r', r'', r^{(3)}, r^{(4)}, z, z', z'', z^{(3)}, z^{(4)}; \kappa, \sigma, p] = 0 \tag{S5}$$

and

$$eqz[r, r', r'', , r^{(3)}, r^{(4)}, z, z', z'', z^{(3)}, z^{(4)}; \kappa, \sigma, p] = 0. \tag{S6}$$

Here we do not show the explicit form of the equations because it is too lengthy and does not contain much information. However, we stress that both equations are fourth-order ordinary differential equations. The solutions to these two equations are not unique because there is a freedom to choose the scaled factor. To see this, note that $[r(u), z(u)]$ and $[r(u^2), z(u^2)]$ give the same membrane shape when $u$ is varied from 0 to 1. In fact, $u^2$ in the bracket can be replaced with any monotonous function $g(u)$ that maps the interval $[0, 1]$ to itself. To avoid the non-uniqueness problem, we add another coordinate constraint

$$a' = 0. \tag{S7}$$

With this constraint, the parameter $u$ is essentially the rescaled arclength.

#### BOUNDARY CONDITIONS

We have a total of 8 boundary conditions:

$$\begin{cases} r(0) &= 0, \\ r(1) &= R_b, \\ r''(0) &= 0, \\ r''(1) &= 0, \\ z(0) &= z_0, \\ z(1) &= 0, \\ z'(0) &= 0, \\ z'(1) &= 0. \end{cases} \tag{S8}$$

The first line is due to axisymmetry of the membrane shape. However, for numerical calculations, $r(0) = 0$ introduces singularities as it appears in the denominator of the curvature expressions. In practice, we choose $r(0) = 0.001$, which is a very small number such that it would not influence the accuracy of the solution at the rest of the points. The second line sets the base radius $R_\mathrm{b}$, which would be learned by the neuron network in the inverse problem. The third line is also due to axisymmetry of the membrane shape. The fifth and the sixth lines set the membrane heights. In the inverse problem, we set $z_0$ to be the same as the experimental profile. The seventh and eighth lines set the angle at the tip and at the base to be zero.

As for the fifth line, it arises from the coordinates constraint (S7) expressed at the boundary

$$a'(1) \propto r'(1)r''(1) + z'(1)z''(1) = r'(1)r''(1) = 0. \tag{S9}$$

The fact that $r'(1)$ is not equal to zero leads to $r''(1) = 0$.

#### LOSS FUNCTION OF THE FORWARD PROBLEM

The forward problem is equivalent to solve the variational equations (S5) and (S6) with the coordinates constraint (S7) and the boundary conditions (S8). Therefore, the loss function of the forward problem $L_\mathrm{for}$ can be written down as the sum of three terms

$$L_\mathrm{for} = L_\mathrm{eqs} + L_\mathrm{con} + L_\mathrm{bc}, \tag{S10}$$

including $L_\mathrm{eqs}$ from the two variational equations, $L_\mathrm{con}$ from the coordinates constraint, and $L_\mathrm{bc}$ from the boundary conditions. Their explicit expressions are given below:

$$L_\mathrm{eqs} = \frac{1}{N+1}\sum_{i=1}^{N-1} eqr(i)^2 + \frac{1}{N+1}\sum_{i=1}^{N-1} eqz(i)^2 \tag{S11}$$

$$L_\mathrm{con} = \frac{1}{N+1}\sum_{i=1}^{N-1} a'(i)^2 \tag{S12}$$

$$\begin{aligned} L_\mathrm{bc} &= [r(0) - 0.001]^2 + [r(1) - R_b]^2 + r''(0)^2 + r''(1)^2 \\ &+ [z(0) - z_0]^2 + z'(0)^2 + z'(1)^2 + z(0)^2 \end{aligned} \tag{S13}$$

where $f(i)$ represents the value of the function $f$ evaluated at $u = \frac{i}{N}$. When the expression involves derivatives, we always use central difference scheme for the middle points, and forward/backward difference scheme for the boundary points. The total number of points is $N + 1$, with the first point at $u = 0$ and the last point at $u = 1$. We find that $N = 16$ is able to solve the equations to a good accuracy in a reasonable time.

We are now in a position to train the neuron network to minimize the loss function $L_\mathrm{for}$. Based on our experiences, we find that applying a so-called Hard Constraints Method (2) has a higher prediction accuracy than directly minimizing the loss function $L_\mathrm{for}$ (Eq. S10). The idea of the Hard Constraints Method is to introduce a transformation $r(u) \to \widetilde{r}(u)$, and $z(u) \to \widetilde{z}(u)$ such that the transformed $\widetilde{r}(u)$ and $\widetilde{r}(u)$ automatically satisfy the boundary conditions. Specifically, note that there are 4 boundary conditions for each neural network output $r(u)$ and $z(u)$, we introduce the following transformations

$$\widetilde{r}(u) = r(u) + a_1 + b_1 u + c_1 u^2 + d_1 u^3, \tag{S14}$$

and

$$\widetilde{z}(u) = z(u) + a_2 + b_2 u + c_2 u^2 + d_2 u^3, \tag{S15}$$

and the coefficients in the polynomials can be solved by requiring $\widetilde{r}(u)$ and $\widetilde{z}(u)$ to fullfil the boundary conditions

$$\begin{cases} \widetilde{r}(0) &= 0.001 &= r(0) + a_1 \\ \widetilde{r}(1) &= R_b &= r(1) + a_1 + b_1 + c_1 + d_1 \\ \widetilde{r}''(0) &= 0 &= r''(0) + a_1 + b_1 + c_1 \\ \widetilde{r}''(1) &= 0 &= r''(1) + a_1 + b_1 + c_1 + d_1 \end{cases} \tag{S16}$$

and

$$\begin{cases} \widetilde{z}(0) &= z_0 &= z(0) + a_2 \\ \widetilde{z}(1) &= 0 &= z(1) + a_2 + b_2 + c_2 + d_2 \\ \widetilde{z}'(0) &= 0 &= z'(0) + b_2 \\ \widetilde{z}'(1) &= 0 &= z'(1) + b_2 + 2c_2 + 3d_2 \end{cases} . \tag{S17}$$

The solutions of the coefficients are

$$\begin{cases} a_1 &= -\frac{r''(1)}{6} - \frac{b_1}{3} \\ b_1 &= -\frac{r''(0)}{2} \\ c_1 &= R_b - r(1) - a_1 - b_1 - d_1 \\ d_1 &= 0.001 - r(0) \end{cases} \tag{S18}$$

and

$$\begin{cases} a_2 &= z_0 - z(0) \\ b_2 &= -z'(0) \\ c_2 &= -3a_2 - 2b_2 - 3z(1) + z'(1) \\ d_2 &= -a_2 - b_2 - c_2 - z(1) \end{cases} . \tag{S19}$$

After the transformation, we can remove the boundary condition term $L_{\text{bc}}$ in the loss function $L_{\text{for}}$ to get a simpler loss function

$$L_{\text{for}} = \frac{1}{N+1} \sum_{i=1}^{N-1} \widetilde{eqr}(i)^2 + \frac{1}{N+1} \sum_{i=1}^{N-1} \widetilde{eqz}(i)^2 + \frac{1}{N+1} \sum_{i=1}^{N-1} \widetilde{a}'(i)^2, \tag{S20}$$

where $\widetilde{eqr}$, $\widetilde{eqz}$ and $\widetilde{a}'$ become equations with $\widetilde{r}$, $\widetilde{z}$ and their derivatives:

$$\widetilde{eqr} = eqr[\widetilde{r}, \widetilde{r}', \widetilde{r}'', \widetilde{r}^{(3)}, \widetilde{r}^{(4)}, \widetilde{z}, \widetilde{z}', \widetilde{z}'', \widetilde{z}^{(3)}, \widetilde{z}^{(4)}; \kappa, \sigma, p] \tag{S21}$$

$$\widetilde{eqz} = eqz[\widetilde{r}, \widetilde{r}', \widetilde{r}'', \widetilde{r}^{(3)}, \widetilde{r}^{(4)}, \widetilde{z}, \widetilde{z}', \widetilde{z}'', \widetilde{z}^{(3)}, \widetilde{z}^{(4)}; \kappa, \sigma, p]. \tag{S22}$$

#### SYMMETRIZATION ALGORITHM

In order to learn the model parameters, we need to compare the ML output with the experimental data, which is structured in pairs of coordinates $(r_i, z_i)$. One example of the experimental profile is shown in Fig. S1. As our model assumes rotational symmetry of the membrane shapes, a symmetrization procedure is applied on the experimental profile to be compared with the ML output.

The procedure is conducted in two steps: First, we take the axis where the maximum value of $z$ is located as the axis of symmetry, and normalize the membrane shapes on the left and right sides using the rescaled arclength $u$, as illustrated in Fig. S1 a; Second, we average $r$ and $z$ with the same $u$ on the left and right sides to obtain the symmetrized shape, as illustrated in Fig S1 b.

#### LOSS FUNCTION OF THE INVERSE PROBLEM

Given an experimental profile, the inverse problem aims to find model parameters such that the ML output not only satisfies the variational equations, but also matches the symmetrized experimetal profile. The loss function of the inverse problem therefore needs to incorporate $L_{\text{data}}$ which measures the difference between the ML output and the symmetrized experimental profile. Specifically, we do interpolation of $r(z_j)$ at equally spaced $z_j = z_0 + (5 - z_0)\frac{j}{50}$, $j = 1, \cdots, 50$ based on the ML output $(r, z)$ pairs and the symmetrized experimental profile $(r, z)$ pairs, respectively. The loss function $L_{\text{data}}$ then takes the squared distance between the two interpolated datasets, i.e.,

$$L_{\text{data}} = \sum_{j=1}^{50} [r_{\text{ML}}(j) - r_{\text{exp}}(j)]^2, \tag{S23}$$

in which $r_{\mathrm{ML}}(j)$ and $r_{\mathrm{exp}}(j)$ represent the interpolation results of the ML output and the symmetrized experimental profile at the same point $z_j$, respectively. The total loss function of inverse problem then reads

$$L_{\mathrm{tot}} = \frac{1}{N+1}\sum_{i=1}^{N-1}\widetilde{eqr}(i)^2 + \frac{1}{N+1}\sum_{i=1}^{N-1}\widetilde{eqz}(i)^2 + \frac{1}{N+1}\sum_{i=1}^{N-1}\widetilde{a}'(i)^2 + \frac{1}{50}\sum_{j=1}^{50}[r_{ML}(j) - r_{exp}(j)]^2. \quad (S24)$$

#### CALCULATION OF THE FORCE

In order to calculate the force $f$ to pull the membrane to a certain height $z_0$, we adopt another form of the variational equations by introducing a new variable $\psi(u)$, which is the tangent angle spanned between the tangential direction and the horizontal direction (3). The tangential angle $\psi(u)$ satisfies the following geometric relations:

$$r'(u) = a(u)\cos\psi(u), \quad (S25)$$

and

$$z'(u) = -a(u)\sin\psi(u). \quad (S26)$$

We also introduce another term in the free energy

$$E_f = -f[z(0) - z(1)] \quad (S27)$$

to account for the work done by a point force $f$ to pull the membrane from 0 to $z_0$. The force $f$ can be considered as a Lagrangian multiplier to impose the membrane height $z(0) - z(1)$. The total free energy of membrane can be rewritten as

$$L = 2\pi\int \mathcal{L}[r, r', z, z', \psi, \psi', a, \alpha, \beta; \kappa, \sigma, p, f]du, \quad (S28)$$

in which

$$\mathcal{L} = \frac{1}{2}\kappa\left(\frac{\sin\psi}{r} + \frac{\psi'}{a}\right)^2 ra + \sigma ra + \frac{p}{2}r^2 a\sin\psi - \frac{f}{2\pi}a\sin\psi + \alpha(r' - a\cos\psi) + \beta(z' + a\sin\psi). \quad (S29)$$

Here $\alpha(u)$ and $\beta(u)$ are two Lagrangian multipliers to impose the geometric relations (S25) and (S26). By performing variations to the variables $\psi, r, z$, respectively, we obtain 3 corresponding equations

$$\psi'' = \frac{a^2 pr\cos\psi}{2\kappa} - \frac{a^2 f\cos\psi}{2r\kappa\pi} + \frac{a^2\alpha\sin\psi}{r\kappa} + \frac{a^2\cos\psi\sin\psi}{r^2} + \frac{a^2\beta\cos\psi}{r\kappa} - \frac{a\psi'\cos\psi}{r}, \quad (S30)$$

$$\alpha' = a\sigma + apr\sin\psi - \frac{a\kappa(\sin\psi)^2}{2r^2} + \frac{\kappa(\psi')^2}{2a}, \quad (S31)$$

and

$$\beta' = 0. \quad (S32)$$

By performing variations to the variable $a$, we obtain a conserved quantity

$$H(u) = \frac{1}{2}pr^2\sin\psi - f\sin\psi - \alpha\cos\psi + \beta\sin\psi + \frac{1}{2}\kappa r\left[(\frac{\sin\psi}{r})^2 - (\frac{\psi'}{a})^2\right] + \sigma r = 0. \quad (S33)$$

Eqs. (S25), (S26), and (S30), (S31), (S32) constitute a system of ordinary differential equations, in which only Eq. (S30) is second order, and all the others are first order. The system is equivalent to a total number of 6 first order ordinary differential equations. In addition, we have two unknown parameters, one being the scaling constant $a$, the other being the force $f$. To complete the equations, we need a total number of 8 boundary conditions. They are listed below:

$$\begin{cases} \psi(0) & = & 0 \\ r(0) & = & 0 \\ z(0) & = & z_0 \\ \beta(0) & = & 0 \\ \mathcal{H}(0) & = & 0 \\ \psi(1) & = & 0 \\ z(1) & = & 0 \\ r(1) & = & R_b \end{cases}. \quad (S34)$$

Finally, we use the MATLAB solver bvp4c, which is based on a finite difference scheme, to solve the set of ordinary differential equations with boundary conditions. The unknown parameter $f$ can be directly extracted from the solution.

#### TRANSFER LEARNING STRATEGY

Based on our experiences, if the neural network starts directly from a random initialization to learn the solution with a large membrane height, it usually takes more than an hour, and sometimes may not even converge. In order to solve this problem, we adopt a training strategy called Transfer Learning (4). The idea is that we start with a random initialization of the neuron network to train the experimental profiles with low membrane heights, and then use the trained network to be the initialized network for profiles with large membrane heights. Specifically, we sort the 79 experimental profiles from low to high, and index them with their cardinal numbers. We then divide the experimental profiles into 8 groups according to Table S1. For each group, the neuron network starts from a random initialization and the training is ordered from low index to high index, i.e., the trained network for one indexed profile would be used as the initial network for training the next indexed profile.

Table S1: Transfer learning strategy.

| group0 | group1 | ... | group6 | group7 |
|---|---|---|---|---|
| index0 | index1 | ... | index6 | index7 |
| index8 | index9 | ... | index14 | index15 |
| | | ... | | |
| index64 | index65 | ... | index70 | index71 |
| index72 | index73 | ... | index78 | |

#### REFERENCES


1. Rahimi, M., and M. Arroyo, 2012. Shape dynamics, lipid hydrodynamics, and the complex viscoelasticity of bilayer membranes. *Physical review E* 86:011932.

2. Lu, L., R. Pestourie, W. Yao, Z. Wang, F. Verdugo, and S. G. Johnson, 2021. Physics-informed neural networks with hard constraints for inverse design. *SIAM Journal on Scientific Computing* 43:B1105–B1132.

3. Jülicher, F., and U. Seifert, 1994. Shape equations for axisymmetric vesicles: a clarification. *Physical Review E* 49:4728.

4. Pan, S. J., and Q. Yang, 2010. A Survey on Transfer Learning. *IEEE TRANSACTIONS ON KNOWLEDGE AND DATA ENGINEERING* 22:1345–1359.

#### SUPPLEMENTARY FIGURE

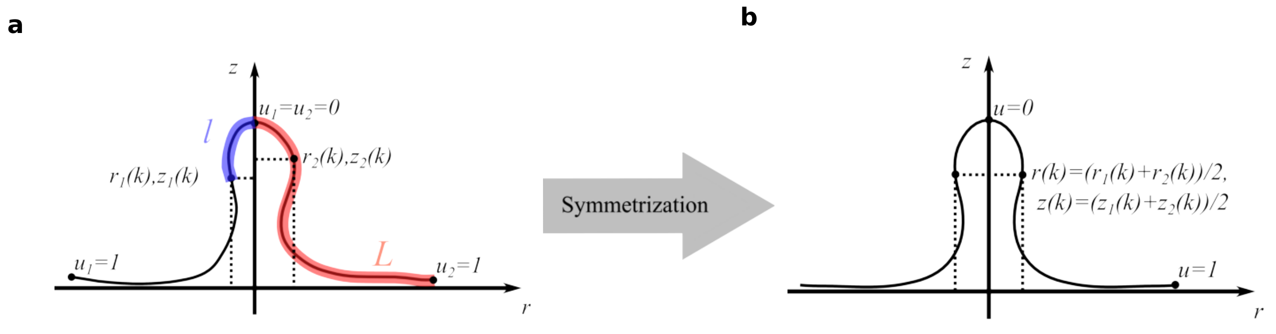


Figure S1: Symmetrization of the experimental profile. (a) The experimental profile curve is divided into a left part and a right part from the highest point. Each part has its rescaled arclength $u_\mathrm{i} = l_\mathrm{i}/L_\mathrm{i}(i = 1, 2)$, where $l_\mathrm{i}$ is the arclength calculated from the highest point, and $L_\mathrm{i}$ is the total arclength to the last point. (b) Symmetrized experimental profile by taking the average of the left part and the right part at the same rescaled arclength.

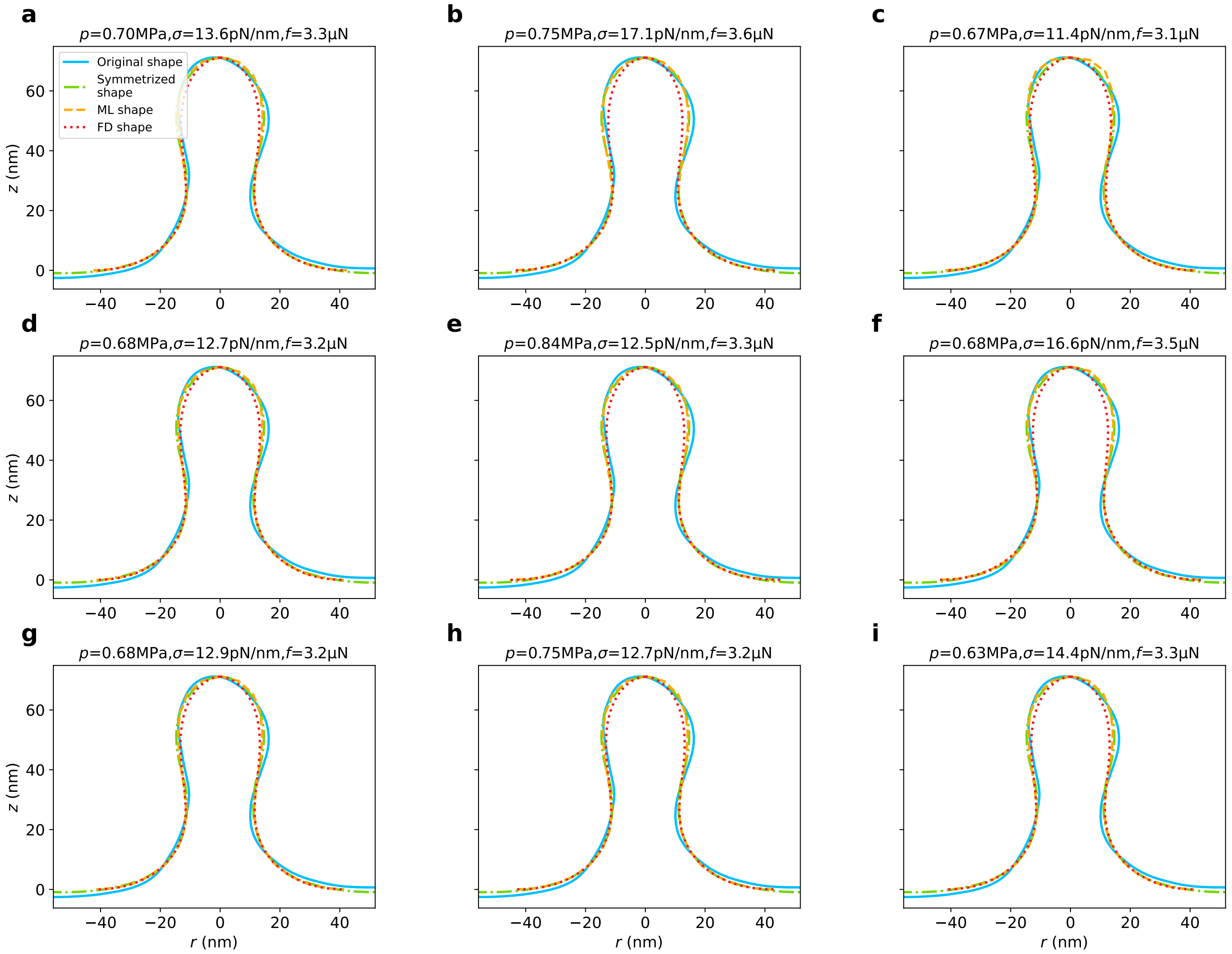


Figure S2: Another 9 learning results for a single experimental membrane profile. The average values of $p$, $\sigma$ and $f$ are 0.71 MPa, 13.7 pN/nm and 3.3 $\mu$N; the stander deviations of $p$, $\sigma$ and $f$ are 0.06 MPa, 0.18 pN/nm and 0.15 $\mu$N.
